# SMARTdb: An Integrated Database for Exploring Single-cell Multi-omics Data of Reproductive Medicine

**DOI:** 10.1101/2023.09.10.556986

**Authors:** Zekai Liu, Zhen Yuan, Yunlei Guo, Ruilin Wang, Yusheng Guan, Zhanglian Wang, Yunan Chen, Tianlu Wang, Meining Jiang, Shuhui Bian

**Affiliations:** State Key Laboratory of Reproductive Medicine and Offspring Health, Nanjing Medical University, Nanjing 211166, China; Collaborative Innovation Center for Cancer Personalized Medicine, School of Public Health, Nanjing Medical University, Nanjing 211166, China

**Keywords:** Database, Single-cell multi-omics, Embryo, Germ cell, Gonad

## Abstract

Single-cell multi-omics sequencing has greatly accelerated reproductive research in recent years, and the data are continually growing. However, utilizing these data resources is challenging for wet-lab researchers. A comprehensive platform for exploring single-cell multi-omics data related to reproduction is urgently needed. Here we introduce SMARTdb (single-cell multi-omics atlas of reproduction), which is an integrative and user-friendly platform for exploring molecular dynamics of reproductive development, aging, and disease, covering multi-omics, multi-species, and multi-stage data. We have curated and analyzed single-cell transcriptome and epigenome data of over 2.0 million cells from 6 species across whole lifespan. A series of powerful functionalities are provided, such as “Query gene expression”, “DIY expression plot”, “DNA methylation plot”, and “Epigenome browser”. With SMARTdb, we found that the male germ-cell-specific expression pattern of *RPL39L* and *RPL10L* is conserved between human and other model animals. Moreover, DNA hypomethylation and open chromatin may regulate the specific expression pattern of *RPL39L* collectively in both male and female germ cells. In summary, SMARTdb is a powerful platform for convenient data mining and gaining novel insights into reproductive development, aging, and disease. SMARTdb is publicly available at https://smart-db.cn.

## Introduction

Single-cell multi-omics technique has greatly promoted our molecular understanding of early embryos [1–6], fetal germ cells and gonadal somatic cells [7–11], developing testis and ovary [12–16], adult testis and ovary [17–26], aging testis and ovary [27–29], endometrium [30,31], maternal-fetal interface [32–34], as well as reproductive diseases [35–39] in the last decade. Multiple molecular layers (including gene expression patterns, DNA methylation dynamics, and open chromatin features) have been profiled with high-throughput sequencing technology, and the data have been growing continually. It is noted that previous single-cell multi-omics studies typically provide comprehensive atlas rather than dig into individual genes or genomic sites, but they offer valuable resources for more wet-lab researchers to perform further investigations into specific questions, genes, and genomic sites of interest. For example, it is important for researchers using model animals to know if the expression pattern of a specific gene is conserved in humans, and whether knocking out of the gene would potentially result in important phenotypes.

However, analyzing the massive and complex single-cell multi-omics data requires experienced skills and a large number of computing resources, which is challenging and time-consuming for wet-lab researchers especially. For instance, as each of those studies typically focuses on a narrow time span, researchers who wish to query the expression levels of a particular gene in child, young adult, aging adult and infertile man need to download and process large amount of data from several separate studies, and they might find this process difficult and laborious due to a lack of analytical skills and computing resources. This inconvenience hinders the optimal utilization of the published data, thus limiting potential novel discoveries. Hence, integrated and user-friendly platforms are needed to bridge the gap between rich data resource and wet-lab researchers.

Endeavors such as human cell atlas (HCA, https://www.humancellatlas.org/) and human protein atlas (HPA, https://www.proteinatlas.org/) have served as excellent platforms for exploring massive single-cell data conveniently. However, these databases do not contain much reproduction-related data so far. Currently, there are a few small-scale reproduction-related websites available, which typically only include data of their own study, are limited in data amount and time span, only provide simple function of querying gene expression of a specific gene, and lack long-term maintenance. As a result, there is still an urgent need for an integrated and user-friendly platform which focuses on promoting efficient utilization of public single-cell multi-omics data related to reproduction across full life-cycle.

Here we introduce SMARTdb (single-cell multi-omics atlas of reproduction), which is a manually curated, integrative and interactive platform aiming to bridge the gap between mass data resources and wet-lab researchers, and promote exploring molecular dynamics of reproductive development, aging, and disease.

## Data collection and processing

### Data collection

We curated single-cell transcriptome and epigenome (DNA methylation and chromatin accessibility) data related to reproduction from published studies (Table S1). The gene expression matrix, DNA methylation, and chromatin accessibility data were downloaded from NCBI GEO (https://www.ncbi.nlm.nih.gov/geo), EMBL-EBI (https://www.ebi.ac.uk/), and Zenodo (https://zenodo.org/). In addition to data of developmental stages and normal physiological states, data of aging and human infertility (such as non-obstructive azoospermia) are also incorporated. In summary, the single-cell data of over 2.0 million individual cells across 6 species (human, monkey, mouse, pig, buffalo, and goat) were curated, involving early embryos, fetal germ cells and gonadal somatic cells, developing testis and ovary, adult testis and ovary, aging testis and ovary, endometrium, maternal-fetal interface, and human infertility. These data cover the entire life cycle, and span > 120 specific time points of 6 species (Table S1).

### Processing of single-cell transcriptome data and cell clustering

To reproduce the cell clustering results of original studies as much as possible, for the datasets with metadata provided by the original study, only the cells with cell type identities were used in downstream analyses. For the datasets without provided metadata, the cells were filtered according to the quality control standards mentioned in the original studies. Seurat [40] (version 4.3.0) was used to perform the following processing steps. Briefly, the unique molecular identifier (UMI) count of each gene for each cell was first divided by the total counts for that cell and multiplied by 10,000 (or author-designated parameters), and was then natural-log transformed. The top highly variable genes among cells were selected with “FindVariableFeatures” function with default or author-designated parameters. The normalized data were then centered and scaled, resulting in that the mean expression of each gene across all cells was zero and the standard deviation was one. Principal component analysis (PCA) was performed on the scaled data with the previously determined highly variable genes used as input. Then the cells were clustered with Louvain algorithm. The uniform manifold approximation and projection (UMAP) was used to visualize the cell clustering in low-dimensional space. For the datasets with obvious batch effects, the anchors of different samples were identified with the “FindIntegrationAnchors” function, and then different samples were integrated to correct batch effects with the “IntegrateData” function. For integration of datasets from different studies, strict quality control standards were used to eliminate potential effects of poor quality cells. Only cells with a gene number > 1000 and a mitochondrial gene ratio < 20% were used in downstream analyses. For the datasets without obvious batch effects, no additional correction was performed.

### Cell type annotation and differentially expressed gene analyses of single-cell transcriptome data

A series of analyses were performed to ensure the accuracy of cell identities for each dataset and integrated datasets. For the datasets with metadata provided by the original study, the cells were annotated correspondingly, and classical marker genes for each cell type were checked to confirm accuracy. For the datasets without metadata, the top differentially expressed genes (DEGs) of each cell cluster were identified using the Wilcoxon rank sum test with the “FindAllMarkers” function in Seurat [40] (version 4.3.0) with default parameters (logfc.threshold = 0.25, min.pct = 0.1, return.thresh = 0.01), and their consistency with classical marker genes for each cell type was carefully checked. For the integrated datasets, the cell identities after integration were further checked with the cell type information identified by each dataset before integration. After these steps, the cell type identities can be annotated accurately.

### Visualization of gene expression levels

First, Seurat objects were converted into “anndata” objects with the R package sceasy (version 0.0.7). Then, the dot plots, heatmaps and violin plots were generated with Scanpy [41] (version 1.9.3). For dot plots and heatmaps, both the relative expression levels with and without standard scaling are provided. The standard scale normalization strategy in Scanpy was performed with parameters “standard_scale = ‘var’”, which standardized the expression level of a given gene between 0 and 1 by subtracting the minimum and dividing each by its maximum.

### Processing of single-cell epigenome data

For the epigenetic data generated by scBS-seq [42] and scCOOL-seq [43] techniques, the processed single-cell DNA methylation and chromatin accessibility data provided by the original study were downloaded from GEO database, and were further processed to standard BigWig format with bedtools (version 2.30.0), UCSC bedGraphToBigWig (version 2.9) and bigWigMerge (version 2) for downstream usage. For the data generated by scBS-seq, only CpG sites were retained. For the data generated by scCOOL-seq, WCG (W = A, T) sites were used for DNA methylation analysis and GCH (H = A, C, T) sites were used for chromatin accessibility analysis. For chromatin accessibility data generated by scATAC-seq, the peak regions of each cell type were mainly collected from the authors of the original study.

Considering the sparse coverage of single-cell epigenome data, we merged single-cell epigenome data according to their cell type identities and stages (*e.g.* 2-cell embryo, 8-week female mitotic germ cells, *etc.*), and sequencing techniques (*e.g.* scBS-seq and scCOOL-seq) using custom scripts. Briefly, for the genomic sites only covered by one cell, they were directly retained in the merged files. For the genomic sites covered by multiple cells in the group, the mean values of covered cells were calculated to represent the final states of the sites. For single-cell DNA methylation data, only the genomic sites with DNA methylation levels < 0.1 or > 0.9 in each individual cell were used in downstream merging. The merged single-cell data of cell groups are more convenient and efficient for downstream exploration.

The DNA methylation “Tanghulu” plot was implemented by a custom Python script to visualize DNA methylation state of each CpG site in a narrow genomic region of interest. For single-cell DNA methylation data, only the genomic sites with values < 0.1 or > 0.9 in each individual cell will be plotted. The DNA methylation level is defined as the percentage of methylated sites in all sites. The WashU epigenome browser [44] was incorporated into SMARTdb for convenient visualization of single-cell DNA methylation levels and chromatin accessibility states.

## Implementation of the database website

SMARTdb was developed utilizing Python 3 in conjunction with the Django framework (https://djangoproject.com). The data housed within SMARTdb is systematically organized and accessible through PostgreSQL (https://www.postgresql.org). The user-friendly front-end interface was created using NG-ZORRO and Angular. The platform is hosted using the Nginx web server (http://nginx.org) and is freely accessible online at https://smart-db.cn. For optimal performance, SMARTdb is best used on major web browsers such as Google Chrome, Microsoft Edge, Safari, and Mozilla Firefox, with JavaScript enabled.

## Database content and usage

### Overall design

SMARTdb is an integrative and interactive platform for exploring single-cell multi-omics data related to reproduction. It was designed to have three main features (**Figure 1** and Figure S1). First, single-cell multi-omics data of over 2.0 million individual cells were manually curated and analyzed (Table S1). Users can access gene expression levels, DNA methylation levels and chromatin accessibility levels at single-cell resolution conveniently. Second, the main stages of full life-cycle are covered, including zygote, early embryo, fetus, infant, child, puberty, adult, and aging. More than 120 specific time points are included in total (Table S1). Besides, data of human infertility are also incorporated. Users can not only explore molecular dynamics during full life-cycle, but also compare normal physiological state with disease. Third, considering cross-species conservation analyses are very important, data across six species are included, including human, non-human primate (monkey), mouse, pig, buffalo, and goat. For example, the researchers using mouse as model animals can find out if the expression pattern of the gene of their interest is conserved in human, which will help select more meaningful research targets.

**Figure 1.**
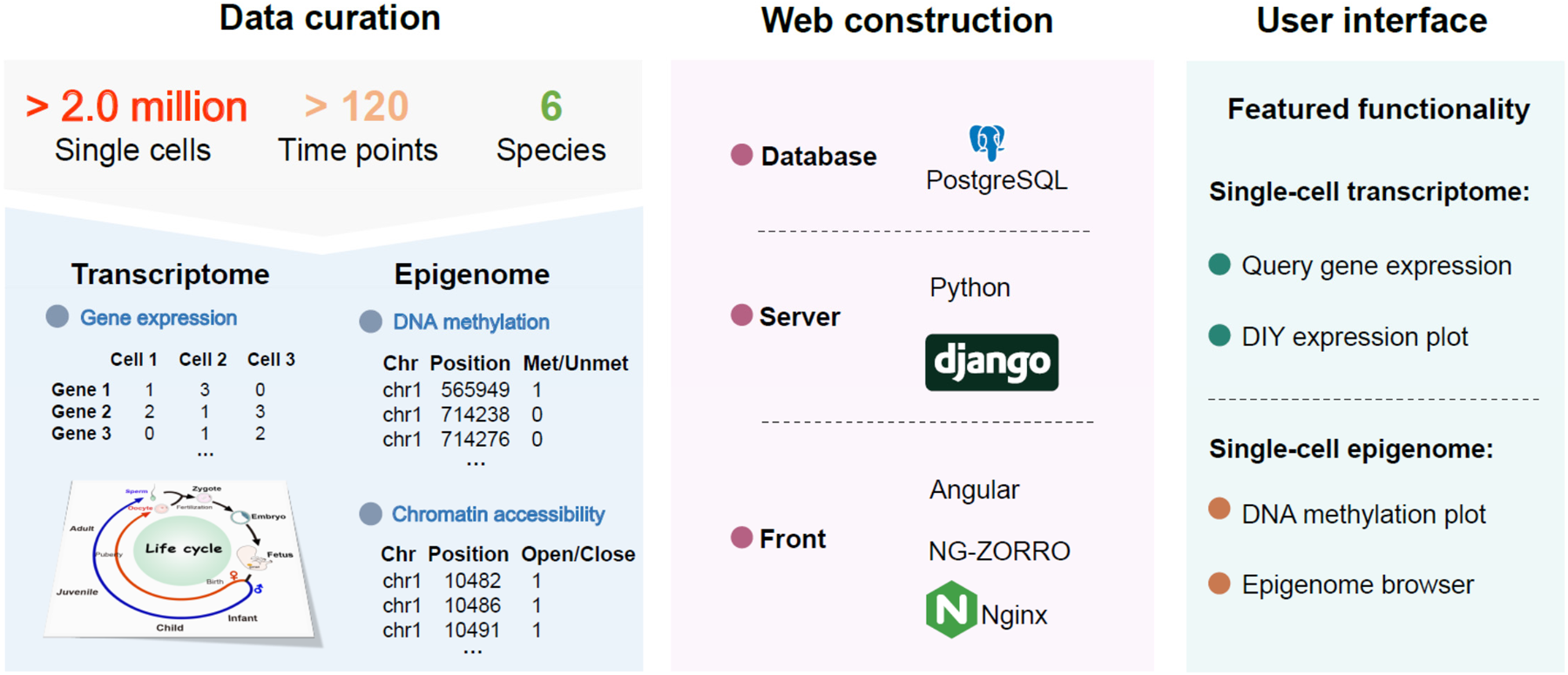
A brief schema describing data curation, web construction, and featured functionalities. The single-cell data of over 2.0 million individual cells across six species (human, monkey, mouse, pig, buffalo, and goat) were curated, covering the entire life cycle and spanning > 120 specific time points of 6 species. Multiple featured function modules were provided, such as “Query gene expression”, “DIY expression plot”, “DNA methylation plot” and “Epigenome browser”.

As a user-friendly platform, SMARTdb offers multiple featured function modules to users for exploring the rich single-cell multi-omics data, such as “Query gene expression”, “DIY expression plot”, “DNA methylation plot” and “Epigenome browser”. The featured function modules are described below.

#### “Overview” and “Query gene expression” modules

In the “Overview” module, SMARTdb facilitates users to get a quick overview of cell clustering, samples, cell counts, and DEGs of each cell type for each scRNA-seq dataset (**Figure 2**A). In the “Query gene expression” module, users can conveniently plot the expression levels of a selected gene across the cell types in the chosen dataset with uniform manifold approximation and projection (UMAP) and violin plots (Figure 2B). Additionally, a concise introduction for each human gene is also provided in this module.

**Figure 2.**
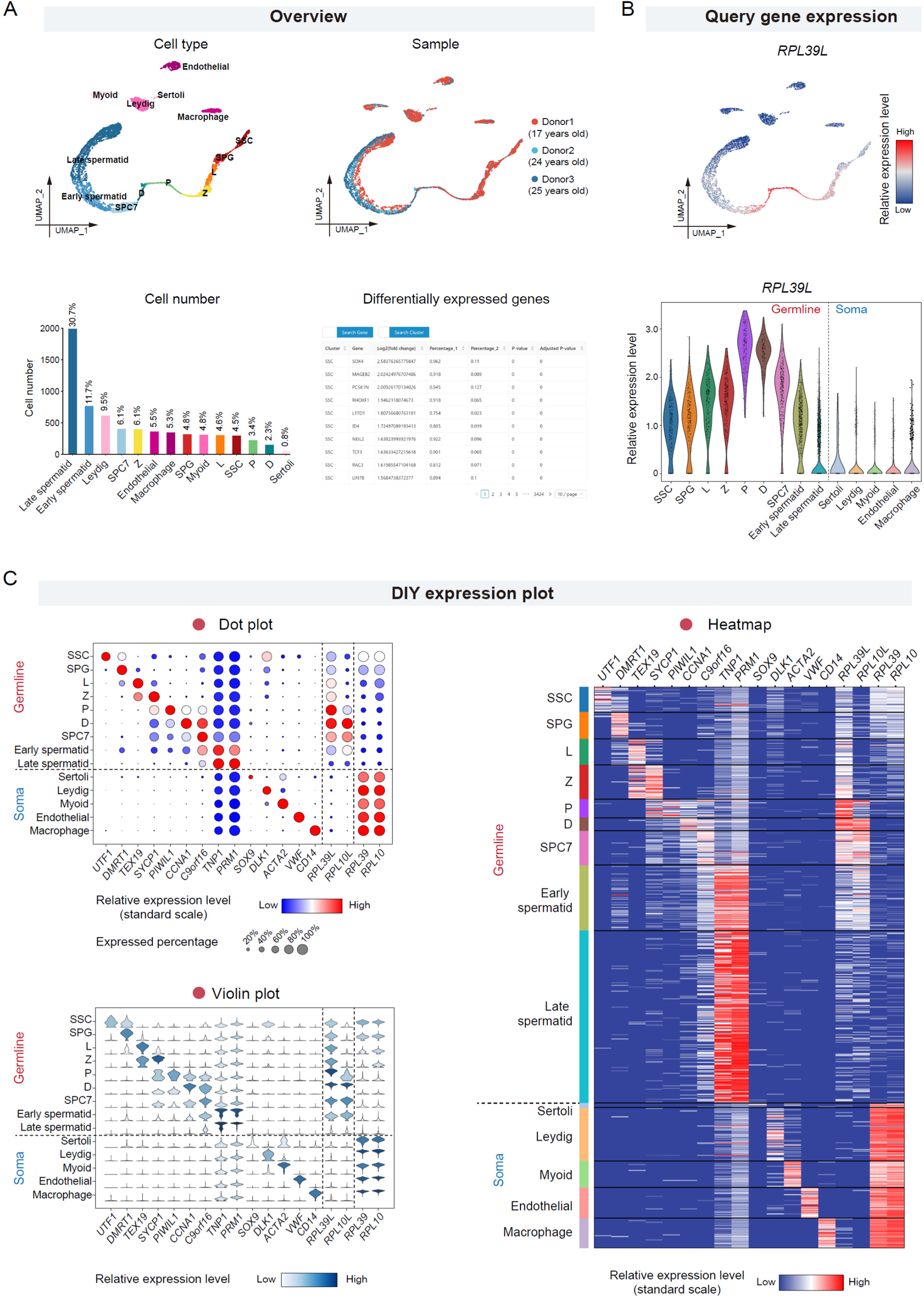
Featured functionalities for exploring single-cell transcriptome datasets. **A.** The “Overview” module provides UMAP plots showing cell types and sample information, bar plots showing cell number and proportion of each cell type, and tables showing differentially expressed genes for each single-cell dataset. The single-cell RNA-seq data of human adult testis (GSE112013) was shown as an example. **B.** The “Query gene expression” module enables visualization of the expression levels of each selected gene with UMAP plot (upper) and violin plot (lower). The expression level of a germ-cell specific ribosomal gene *RPL39L* in human adult testis was shown. **C.** The “DIY expression plot” module allows users to create customized plots displaying the expression levels of selected genes in selected cell types. Three types of plots (dot plot, violin plot and heatmap) are provided. The expression levels of representative marker genes and four ribosomal genes in major cell types of human adult testis were shown. The descriptions of cell type abbreviation are listed below. SSC, spermatogonia stem cell. SPG, spermatogonia. L, leptotene spermatocyte. Z, zygotene spermatocyte. P, pachytene spermatocyte. D, diplotene spermatocyte. SPC7, a mixture of diakinesis, metaphase, anaphase, telophase, and secondary spermatocytes.

#### “DIY expression plot” module

This module provides a very practical function for users to customize expression plots according to their personalized requirements (Figure 2C). Users can select multiple genes and multiple cell types of interest to plot simultaneously, and choose appropriate plot forms for visualization. Three main visualization plots are offered in this module, including dot plots, heatmaps, and violin plots. Moreover, the orders of genes and cell types displayed in the plots can be designated by users freely. Users can also choose whether to standardly scale the expression levels within each gene when using dot plots and heatmaps. The generated figures are of publication quality.

#### “DNA methylation plot” module

This module provides “Tanghulu” plots to visualize DNA methylation status of each cytosine site in CpG or WCG context (**Figure** 3A). Users can conveniently generate a customized “Tanghulu” plot by selecting datasets and cell types, and designating specific genomic region range of interest (such as promoters and gene bodies). In “Tanghulu” plot, each row represents an individual cell, and each circle represents a cytosine site in CpG or WCG context. Black circles stand for methylated sites, while the white circles refer to unmethylated sites. The DNA methylation levels of selected region in each single cell are labeled on “Tanghulu” plots.

**Figure 3.**
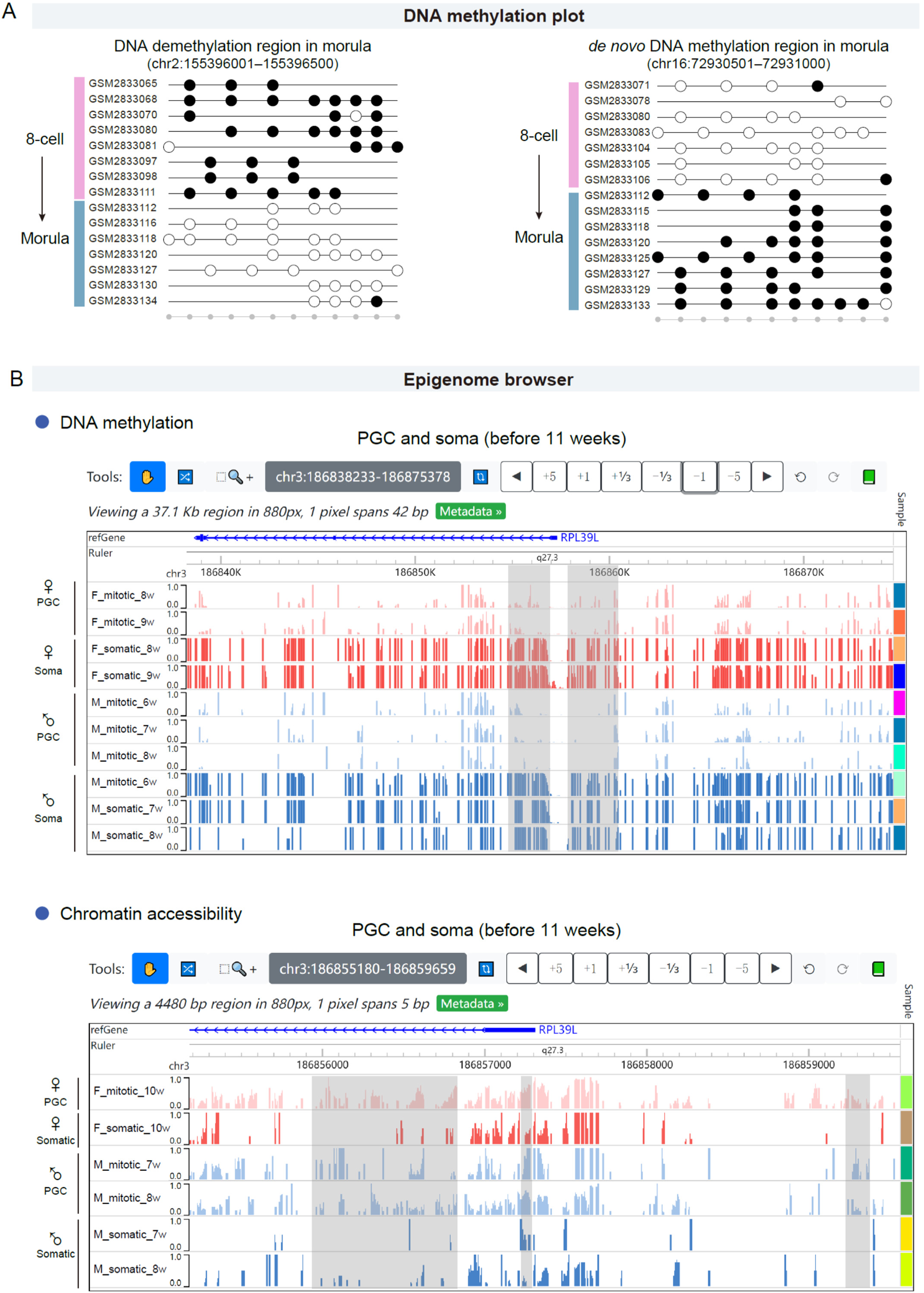
Featured functionalities for exploring single-cell epigenome datasets. **A.** The representative DNA demethylation and *de novo* methylation regions between human 8-cell embryo and morula were shown. Each row in the “Tanghulu” plot corresponds to an individual cell, and each circle represents a CpG site. Methylated CpG sites are represented by black dots, while unmethylated CpG sites are indicated by white dots. The gray dots at the bottom indicate covered CpG sites in selected cells. The CpG sites which were not covered by any cell were omitted. The single-cell DNA methylation data of human early embryo (GSE100272) was used. **B.** The screenshots of epigenome browser displaying the DNA methylation levels and chromatin accessibility levels around the promoter region of *RPL39L* in primordial germ cells. The gray rectangles highlight regions showing significant differences between somatic niche cells and germline cells. PGC, primordial germ cell.

#### “Epigenome browser” module

This module incorporates the WashU epigenome browser tool [44] and offers a state-of-the-art solution for browsing epigenetic datasets (Figure 3B). Users can select single-cell DNA methylation and chromatin accessibility datasets of various cell types based on their interest, such as oocytes, sperms, early embryos, fetal germ cells, and gonadal somatic cells. The DNA methylation and chromatin accessibility levels of selected samples can be directly and conveniently visualized in the browser. Especially, since single-cell data are typically sparse, the single-cell data of each cell type or sample type are merged for optimal visualization and enabling more efficient exploration. The genomic region locator allows users to navigate to a specific gene or designated specific genomic regions (such as promoters and gene bodies), zoom in and out, and move to new regions by dragging the mouse left and right. Additionally, the browser interface is allowed to be customized to meet personalized requirements, such as display mode (bar plot or heatmap), RGB colors, axis scales, and text labels.

### Examples for using SMARTdb

Here we illustrate how to make good use of SMARTdb to facilitate exploration of single-cell multi-omics data and promote new findings. A recent study has revealed that RPL39L is a male germ-cell-specific ribosome protein and plays critical roles in spermatogenesis and male fertility in mouse, whereas the corresponding core ribosome protein RPL39 is mainly expressed in somatic cells [45]. Inspired by the study, we further asked: (1) whether the germ-cell-specific expression pattern is conserved in testis of human, non-human primates as well as other model animals? (2) when does the germ-cell-specific expression pattern of RPL39L form? (3) does epigenetic regulation function in regulating the male germ-cell-specific expression pattern? We will demonstrate how to explore the three questions using SMARTdb below.

#### Cross-species conservation

We utilized the “DIY expression plot” module to plot the expression levels of the ribosomal large subunit protein genes *RPL39L* and *RPL10L*, and the corresponding core ribosome protein genes *RPL39* and *RPL10* in human [23], as well as their homologous genes in mouse [46], monkey [19], and pig [15] (Figures 2C and 4A). For human adult testis [23], we found that the expression levels of *RPL39L* and *RPL10L* peak at spermatocyte, and then reduce in later stages of spermatogenesis at mRNA level (Figure 2B and C). In contrast, *RPL39* and *RPL10* decrease their expression levels during spermatogenesis, and are mainly expressed in somatic cells (Figure 2C). Moreover, the germ-cell-specific expression pattern of *RPL39L* and *RPL10L* is conserved in human and other species (**Figure** 4A).

**Figure 4.**
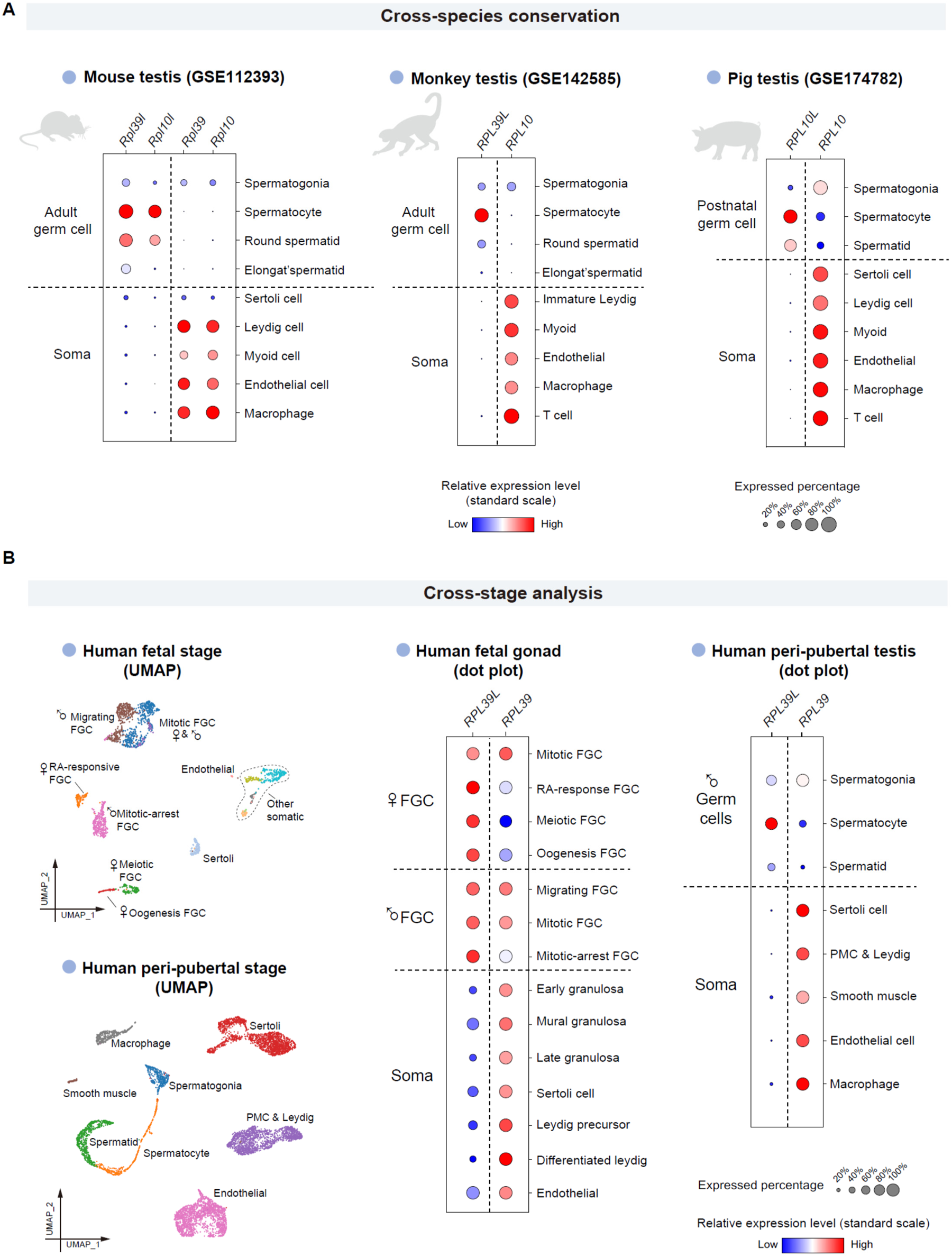
Cross-species and cross-stage analyses of ribosomal genes using SMARTdb. **A.** The dot plots show the expression levels of four ribosomal genes in mouse, cynomolgus monkey, and pig. PMC, peritubular myoid cell. IP, interstitial progenitor cell. **B.** The UMAP plots show the cell type identities of each cell cluster. The expression levels of two ribosomal genes (*RPL39L* and *RPL39*) in human germline and somatic niche cells of fetal stage and peri-pubertal stage were shown in dot plots. FGC, fetal germ cell. PMC, peritubular myoid cell.

#### Developing dynamics

To clarify the second question, we plotted the expression levels of *RPL39L* in fetal stage [10] and peri-pubertal stage [13] with the “DIY expression plot” module (Figure 4B).We found that the germ cells in fetal stages have started to highly express *RPL39L*, as early as in migrating primordial germ cells (PGCs). Moreover, we noticed that *RPL39L* transcripts are highly expressed in both male and female germ cells in human fetal stage, but diverge between male and female in adulthood. In human adulthood, *RPL39L* is relatively highly expressed in male germ cells and female growing oocytes, but lowly expressed in fully-grown, metaphase I (MI), and metaphase II (MII) oocytes [21] (**Figure** 5A). The divergence in the *RPL39L* expression among oocytes of various stages in human adult ovary was detected in mRNA level, indicating potential transcriptional regulation of the gene.

**Figure 5.**
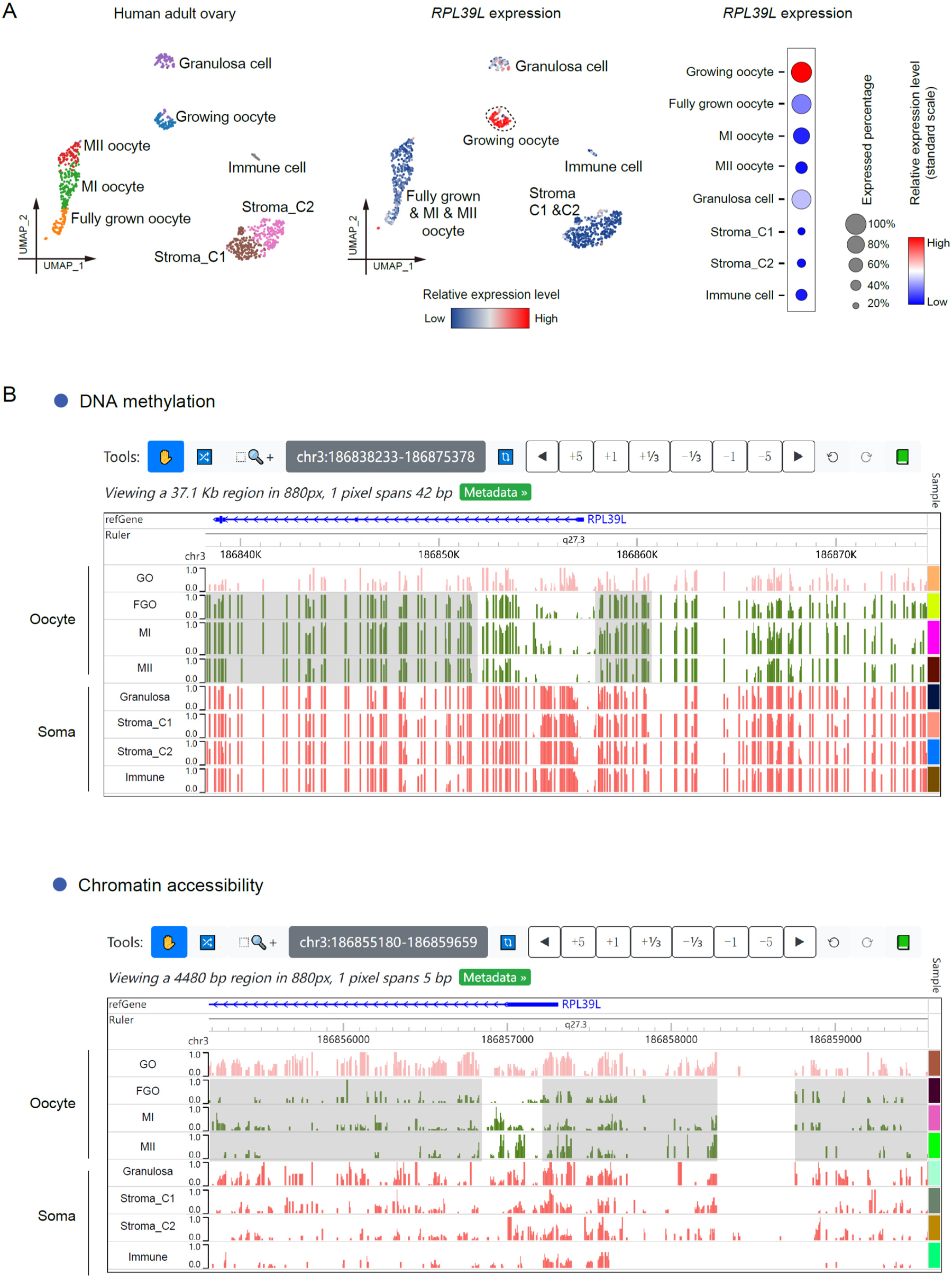
The gene expression, DNA methylation and chromatin accessibility patterns of *RPL39L* in human ovary. **A.** The leftmost UMAP plot shows the cell type identities of human adult ovary. The expression levels of *RPL39L* in various cell types of human adult ovary are shown on the middle UMAP plot and the rightmost dot plot. **B.** The screenshots of epigenome browser displaying the DNA methylation levels and chromatin accessibility levels around the promoter region of *RPL39L* in human adult ovary. The gray rectangles highlight regions showing changes in fully grown, MI, and MII oocytes compared with growing oocytes. GO, growing oocyte. FGO, fully grown oocyte. MI, metaphase I. MII, metaphase II.

#### Regulation mechanism

Then we further explored the transcriptional regulatory mechanism using the “Epigenome browser” module. With epigenetic data of human fetal stage [7], we found that germ cells in fetal stage have hypomethylated and more accessible regions around the promoter of *RPL39L* compared with somatic cells (Figure 3B and Figure S2), which is consistent with the relatively high expression of *RPL39L* of fetal germ cells (Figure 4B). For human adult female [21], growing oocytes displayed similar DNA methylation and chromatin accessibility pattern with fetal germ cells (Figure 5B). In contrast, fully-grown, MI, and MII oocytes up-regulate the DNA methylation levels around *RPL39L* promoter, and down-regulate the chromatin accessibility levels around *RPL39L* promoter (Figure 5B), which may explain the decreased expression level of *RPL39L* in fully-grown, MI, and MII oocytes. Hence, epigenetic regulation potentially plays important roles in shaping the expression pattern of *RPL39L* in both male and female human germ cells during both fetal and adult stages.

## Discussion

The mystery of reproduction-related development, aging and disease has long been explored. Single-cell multi-omics sequencing technique has revolutionized researches in reproduction in recent years, and has led to accelerated accumulation of massive datasets. The public data provide valuable resources for researchers to perform further data mining. However, difficulties in processing big data for broad wet-lab researchers prevent them from being effectively utilized. To deal with the situation of growing data and promote a more efficient usage, we present SMARTdb – a single-cell multi-omics platform for reproductive research. To the best of our knowledge, SMARTdb is the first integrated platform with multiple powerful functionalities to explore single-cell multi-omics data in the field of reproductive medicine. Moreover, it will be continuously updated by adding high-quality single-cell multi-omics data.

Our database will promote reproductive medicine research in several aspects. For example, researchers using model animals can speculate whether knocking out of a specific gene could potentially lead to important phenotypic changes. Users are able to know whether the expression pattern of a specific gene is conserved in human, which will help select more meaningful research targets. For researcher using human samples, cross-species conservation analyses provide important clues for selecting suitable model animals. Cross-stage dynamic analysis will help select the right time window for further investigations. Moreover, epigenetics data resources will provide novel insights for potential regulatory mechanisms.

We showed examples for using SMARTdb and demonstrated the power of it by exploring germ-cell-specific genes. We not only validated the germ-cell-specific expression of ribosome gene *RPL39L*, but also gained a deeper understanding of its cross-species conservation, cross-stage expression dynamics, and potential epigenetic regulation. Moreover, taking the advantages of single-cell transcriptome sequencing, the gene expression patterns in multiple cell types can be observed more accurately and in detail. For example, we found that *RPL39L* is not only specifically expressed in male germ cells in human adult testis, but also highly expressed in the growing oocytes (one subtype of oocytes) in human adult ovary. Furthermore, DNA methylation and chromatin accessibility may cooperate in regulating the divergence of *RPL39L* expression in germ cells of human adult at mRNA level. The results demonstrate the power of SMARTdb, providing valuable clues for further investigations.

In summary, we firmly believe that with the assistance of SMARTdb, researchers in the field of reproduction can easily access and explore massive single-cell multi-omics data, and hopefully generate novel ideas and new findings.

## Data availability

SMARTdb is publicly available at https://smart-db.cn.

## CRediT author statement

**Shuhui Bian**: Conceptualization, supervision, Writing – original draft, Writing – review & editing, Visualization, Investigation, Data curation, Methodology, Formal analysis, Funding acquisition, Project administration. **Zekai Liu:** Investigation, Data curation, Formal analysis, Methodology, Software, Visualization, Validation. **Zhen Yuan:** Investigation, Data curation, Formal analysis, Methodology, Software, Visualization, Validation, Writing – original draft. **Yunlei Guo:** Investigation, Formal analysis, Methodology, Software, Visualization, Validation. **Ruilin Wang:** Investigation, Formal analysis, Validation. **Yusheng Guan:** Investigation, Data curation, Methodology, Validation. **Zhanglian Wang**: Investigation, Data curation. **Yunan Chen**: Investigation, Data curation. **Tianlu Wang**: Investigation, Data curation. **Meining Jiang:** Investigation, Data curation.

## Acknowledgments

We thank professors Fuchou Tang (Peking University), Jingchu Luo (Peking University), Zhibin Hu (Nanjing Medical University), Jiahao Sha (Nanjing Medical University), Fan Guo (Institute of Zoology, Chinese Academy of Sciences), Ran Huo (Nanjing Medical University), and Boqiang Hu (Zhejiang University) for discussions and suggestions. This work was supported by the Young Elite Scientists Sponsorship Program by China Association for Science and Technology (CAST) (Grant No. 2020QNRC001) and research start-up funding from Nanjing Medical University, China (Grant No. KY116RC20200007). All authors read and approved the final manuscript.

## Supplementary material

**Figure S1.**
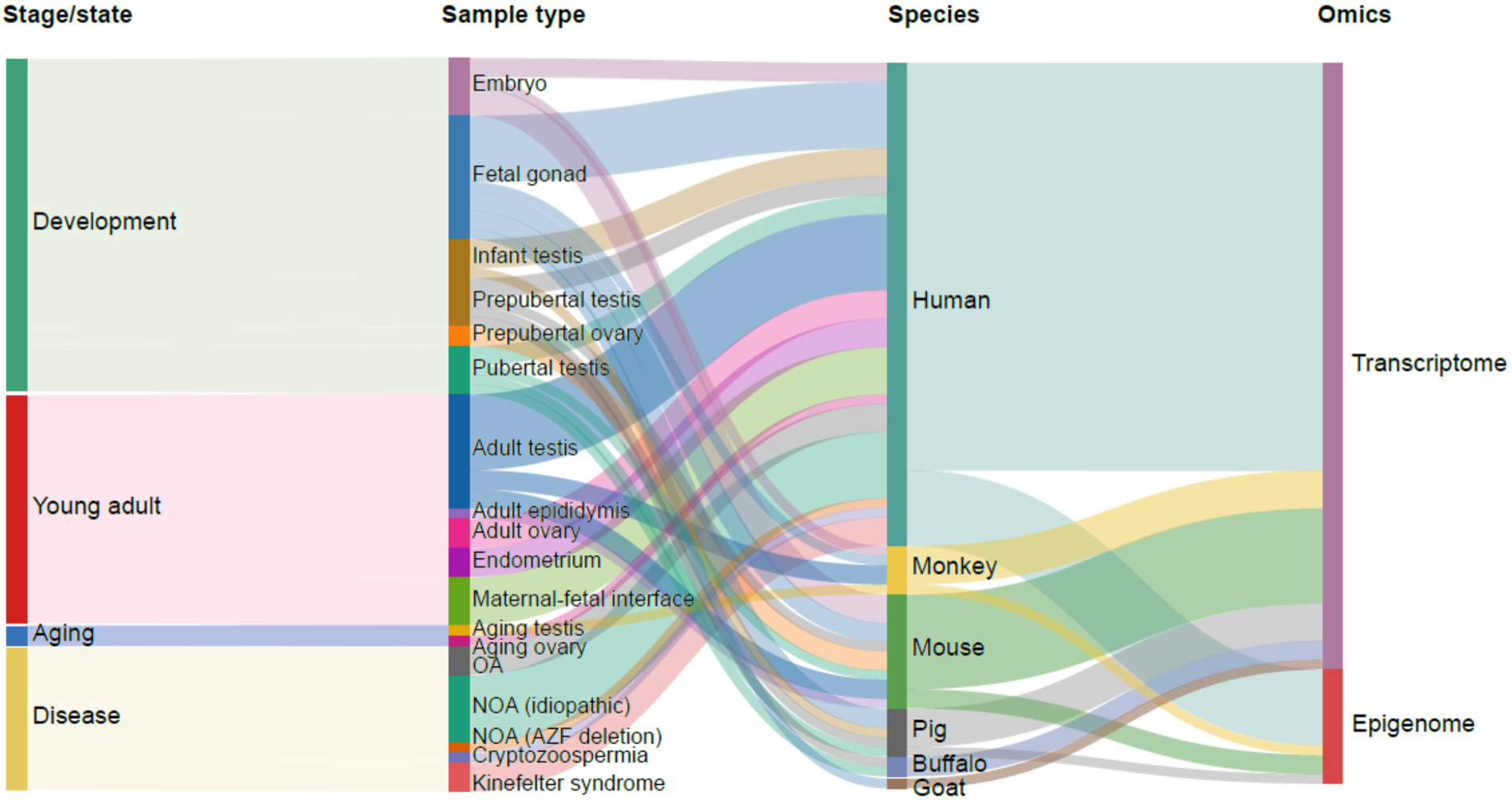
The Sankey plot shows the composition of datasets included in SMARTdb. OA, obstructive azoospermia. NOA, Non-obstructive azoospermia.

**Figure S2.**
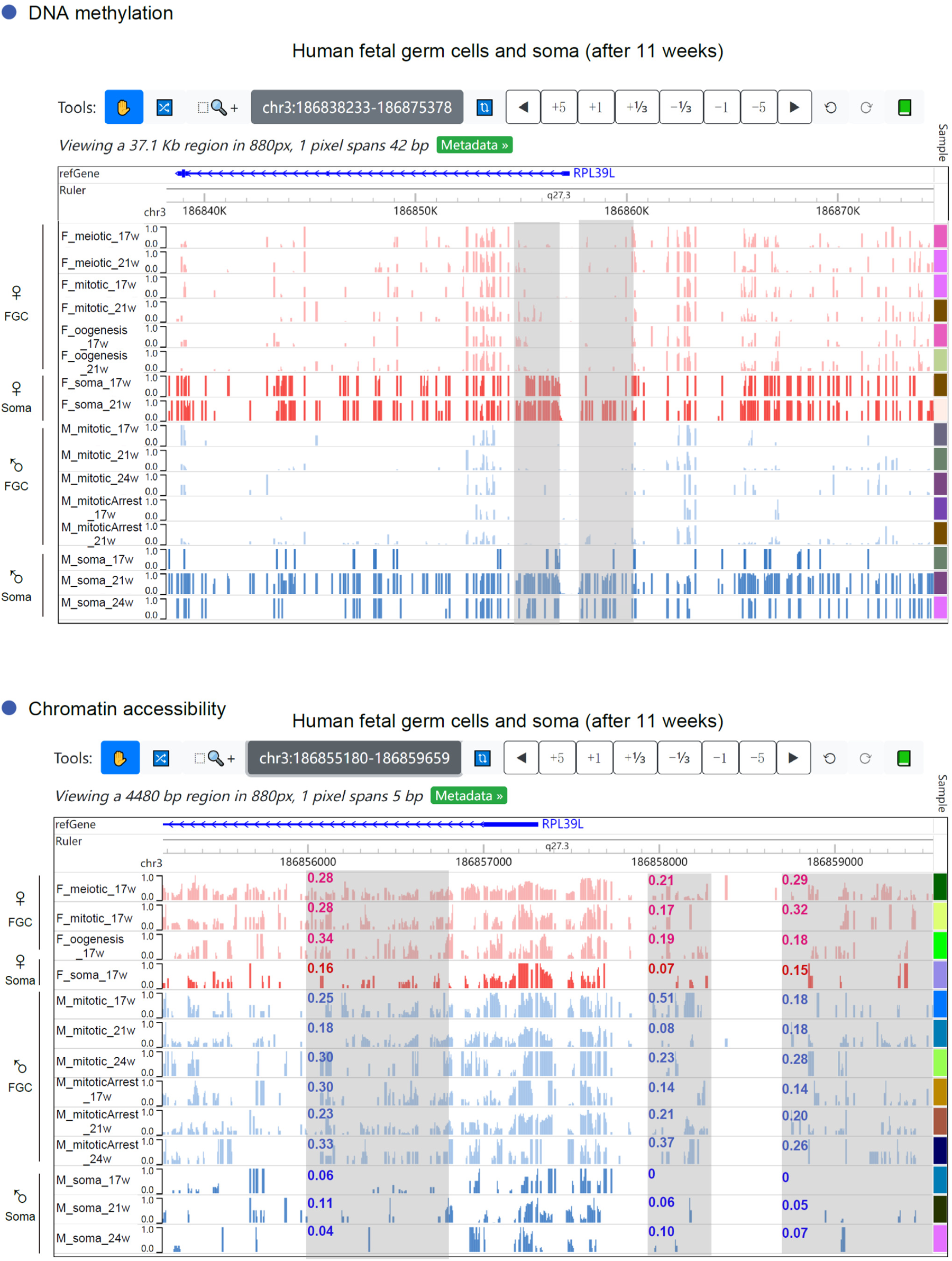
The DNA methylation levels and chromatin accessibility levels around the *RPL39L* promoter in fetal germ cells after 11 weeks. The screenshots of epigenome browser displaying the DNA methylation levels and chromatin accessibility levels around the *RPL39L* promoter in fetal germ cells after 11 weeks post fertilization. The gray rectangles highlight regions showing significant differences between germ cells and somatic niche cells. The mean chromatin accessibility levels of the regions which are highlighted by gray rectangles are labeled.

